# Identifying intervention strategies from machine learning models with COALA: a counterfactual optimization framework

**DOI:** 10.1101/2025.07.18.664723

**Authors:** Bryant Han, Qingling Duan, Ting Hu

## Abstract

**Motivation:** Machine learning (ML) models have become increasingly complex, often functioning as black boxes that limit our understanding of the features contributing to ML predictions. Common explainable AI (XAI) methods such as SHapley Additive exPlanations (SHAP) focus on feature importance but fall short in identifying interactions among features and informing targeted, personalized interventions. Counterfactuals are hypothetical events where specific variables are altered to cause a change in outcome. These causal statements can be applied to AI models to identify actionable interventions for different subjects in a population.

**Results:** We propose the framework Counterfactual Optimization for Actionable interpretabiLity in AI (COALA). COALA interprets models by identifying optimal counterfactuals for each subject, which are defined as actionable changes that lead to the most positive change in predicted outcome. When applied to a gradient boosted tree model trained on the National Health and Nutrition Examination Survey (NHANES) dataset, COALA identifies different profiles of optimal counterfactuals across subjects. Features that remain constrained were able to predict the optimal counterfactual changes for a subject at 85.4%, revealing specific features that drive what a subject’s optimal counterfactual is.

**Availability and Implementation:** Code for COALA implementation, synthetic data, models trained on synthetic data, and code to replicate results and figures are available at https://github.com/brt-solo/COALA. The NHANES 2017–2018 dataset is publicly available from the National Center for Health Statistics (NCHS). The Framingham Heart Study dataset used in this study is a publicly available, Framingham-derived dataset distributed through the Massachusetts Institute of Technology OpenCourseWare (MIT OCW) repository.

## Introduction

### Explainable artificial intelligence

One approach through which artificial intelligence (AI) has advanced the biomedical field is the development of complex machine learning (ML) models to predict health outcomes. Most diseases or clinical phenotypes are complex, with numerous genetic and environmental factors that influence the outcome [Virolainen et al., 2023, Bookman et al., 2011]. The effects of these factors may be additive, such that they independently influence disease risk, or interactive, where the effect of one factor depends on the presence of another factor.

ML models have become increasingly utilized in biomedical research to study complex diseases, using biological factors as features for ML predictions [Xu and Jackson, 2019]. Although ML models have the potential to capture the complexity of biological data, increased model complexity comes with the tradeoff of lower transparency to how features drive predictions [Tanyel et al., 2023, Štrumbelj and Kononenko, 2014, Gunning et al., 2019]. This limitation has prevented the wide-spread application of artificial intelligence to generate novel hypotheses of the biological mechanisms underlying diseases, which may inform prevention and intervention strategies. Even if an ML model can accurately predict a clinical outcome, our understanding of the underlying biology remains limited. The field of explainable AI (XAI) aims to address this gap by identifying key features contributing to prediction models which could improve our understanding of the biological mechanisms contributing to disease outcomes [Gunning et al., 2019, A. and R., 2023].

The most common XAI methods in biomedicine are attribution based, such as Locally Interpretable Model-Agnostic Explainer (LIME) and Shapley Additive exPlanations (SHAP), which assign scores to each feature’s contribution to an individual prediction [Ribeiro et al., 2016, Lundberg and Lee, 2017]. The scores from feature attribution can then be aggregated to produce a feature importance score for each feature [Molnar, 2025]. While feature attribution scores are useful in identifying which variables influence the present predicted outcome, they do not reveal how changing the variables would alter the predicted outcome. To address the limitations of feature attribution, we turn to an XAI approach of counterfactuals [Byrne, 2019].

### Counterfactuals

Counterfactuals are hypothetical events modified from original events that would lead to a different outcome. A counterfactual statement would follow “If X variable is changed by Y, the outcome changes by Z” (e.g., if dietary protein intake is increased, the predicted diabetes risk decreases). This creates a direct link between a variable and the outcome, showing that changing X variable causes a change in the predicted outcome. Existing counterfactual approaches, such as Diverse Counterfactual Explanations (DiCE) [Mothilal et al., 2020] and Model-Agnostic Counterfactual Explanation (MACE) [Yang et al., 2022], can succesfully generate counterfactuals for individual predictions to identify actionable changes that improve the predicted outcome. These counterfactuals may then be used to inform real-life decisions or assess whether the model has learned meaningful relationships in the data [Wielopolski et al., 2024].

Counterfactual tools have been applied in fields where ML models have high performance, such as in financial risk analysis [Wielopolski et al., 2024]. However, there are certain challenges when applying counterfactuals to ML in biomedicine. First, ML models in biomedicine often yield modest performance due to the complexity of biological outcomes and high dimensionality, individual counterfactual explanations may be unreliable in isolation [Beam and Kohane, 2018, Papenmeier et al., 2019]. Identifying consistent trends across a population provides more robust explanations, motivating a population-level counterfactual framework. Second, biological factors can be interconnected in how they affect complex diseases [Virolainen et al., 2023]. Thus, counterfactuals should be generated by changing multiple features to understand how features cumulatively change the predicted outcome. Third, there may be multiple counterfactuals that could result in the same predicted outcome. As such, there is often a seemingly limitless number of possible counterfactuals [Brughmans et al., 2024]. Without a principled selection criterion, counterfactuals are difficult to compare across subjects. There needs to be an objective of identifying a singular optimal counterfactual for each subject, so that counterfactuals may be compared between subjects in a population.

Personalized treatments and interventions are an application of optimal counterfactuals, where different subjects have different optimal changes that improve their health. Interventions may be personalized based on factors that determine the optimal counterfactual, including current health status, previous health history, and family history. We propose a counterfactual framework that identifies optimal counterfactuals across a population and reveals which constraint features determine what the optimal counterfactual is for a given subject, enabling personalized intervention strategies.

### Evolutionary algorithms

To optimize for counterfactuals that lead to the best outcomes, we use evolutionary algorithms, which are search algorithms inspired by biological evolution and natural selection [Holland, 1992]. The counterfactual search space can be high-dimensional and non-differentiable with respect to the input features, making exhaustive search methods and gradient-based optimization unsuitable. Evolutionary algorithms are particularly suited for large, non-convex search spaces where multiple distinct high-fitness solutions may exist simultaneously, as is common in biological optimization problems [Sudholt, 2018].

Evolutionary algorithms have three main characteristics that operate similarly to natural evolution: a population of candidate solutions, fitness, and variation [Holland, 1992, Yu and Gen, 2010, Lutton et al., 2016]. Evolutionary algorithms begin by generating a population of candidate solutions. We define a candidate as a generated counterfactual. Fitness, which is the metric for selection, is defined as the value of the ML model-predicted outcome. Variation refers to operations applied to candidates to generate new candidate solutions, such as through mutation and crossover. Mutation perturbs one or more values of a candidate solution, while crossover combines values from two candidates to produce a new one.

### Framework overview

Building on these principles, we introduce Counterfactual Optimization for Actionable interpretabiLity in AI (COALA), a novel framework that, given a population of subjects, searches for optimal counterfactuals to provide interpretable explanations of the model and potential actionable interventions (Figure 1). COALA draws from the MAP-Elites algorithm, a diversity-focused evolutionary algorithm that partitions the solution space into cells and identifies the optimal solution within each, enabling simultaneous exploration of diverse high-quality solutions [Mouret and Clune, 2015]. COALA partitions the counterfactual search space into cells that are defined by which features are mutable to generate the counterfactual and finds the optimal counterfactual in each cell. COALA allows users to define one or more cells by specifying which feature categories are mutable. In this paper, we primarily apply COALA using a single cell in which real-life actionable features (e.g., dietary intake) are mutable. Multi-cell analyses, which reveal how optimal interventions differ depending on which feature categories are available to change, are demonstrated in the Supplementary Materials.

**Figure 1.**
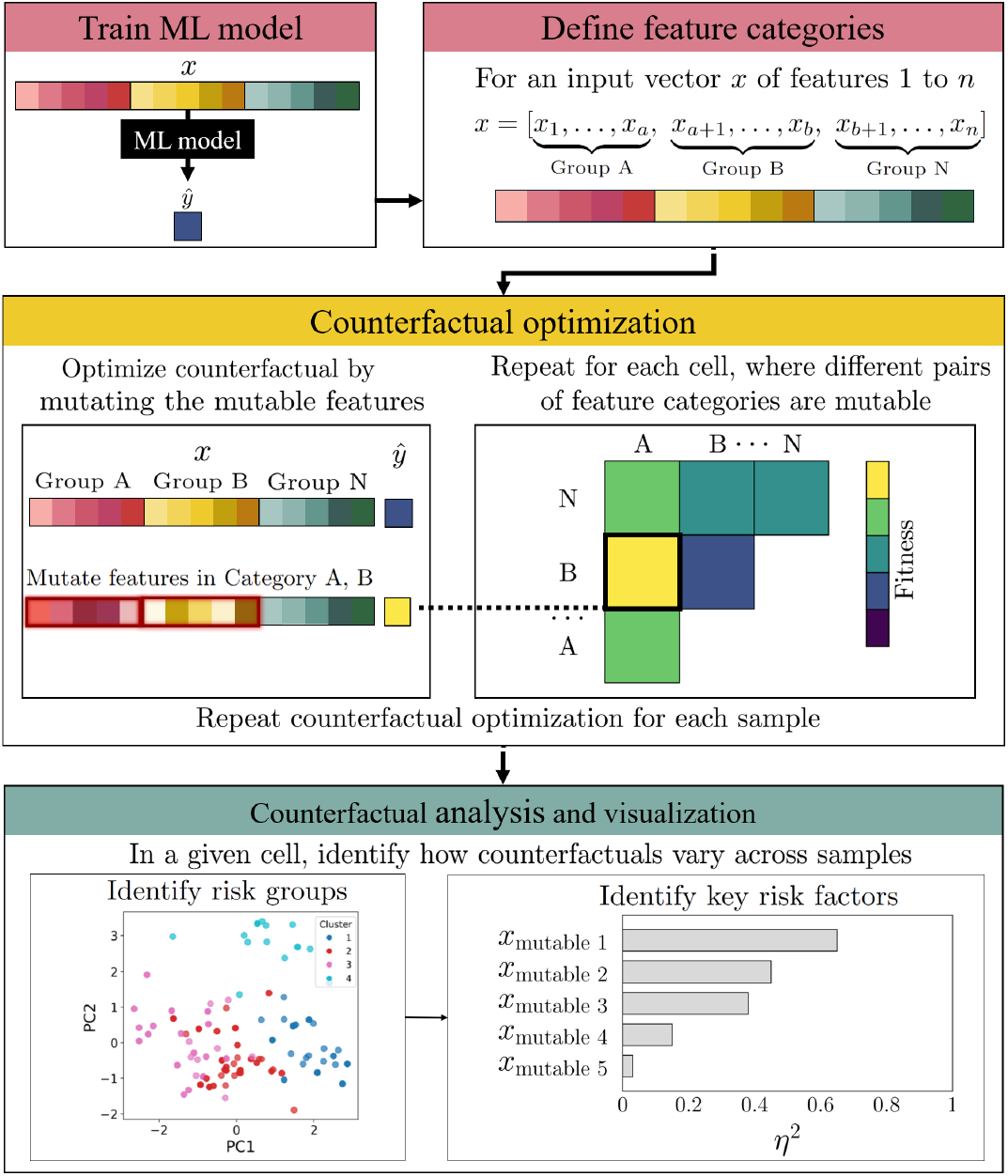
Overview of the pipeline for analyzing counterfactuals with COALA. Values of specific feature categories are changed for an input vector, with high fitness candidates placed in its corresponding cell. For regression models, the fitness is the ML model-predicted outcome. For binary classification, the fitness is the model-predicted probability of the outcome or class. Optimal counterfactuals are expected to vary if there are interactions between constraint and mutable features, such that identified clusters represent groups of patients and their appropriate intervention.

## Methods

### Counterfactual optimization framework

The user first supplies a pre-trained ML model for a prediction task (e.g., predicting disease risk), which is used to evaluate candidate counterfactual solutions based on their predicted outcomes. Next, the user defines feature categories (lines 2–4 of Algorithm 1), which serve to partition the search space into interpretable cells. Each cell represents a combination of feature pairs that could be changed together to create a counterfactual. These feature categories can be specified based on domain-specific knowledge. For instance, in a model trained on electronic health records (EHR), features might be grouped into categories such as diagnosis, demographics, and medication use. In a multi-omics context, each omic layer (e.g., transcriptomics, proteomics, metabolomics) could form a separate category.

A candidate is defined as a generated counterfactual, where the fitness of that candidate is defined as the predicted outcome and prediction confidence score for regression and classification models, respectively. An optimal counterfactual is the candidate with the highest fitness. For each input sample *x*, COALA algorithmically identifies an optimal counterfactual for each cell. Using the input sample as a fixed reference point, COALA begins with a random initialization phase (lines 7–10 and 16–20 of Algorithm 1), where a diverse set of counterfactuals is generated. In each iteration of this phase, a cell (*C*_*i,j*_) is selected at random, and a counterfactual is generated by randomly perturbing the features in the corresponding category pair *F*_*i,j*_. In our basic implementation of COALA, each continuous mutable feature *x*_*k*_ in the feature set *F*_*i,j*_ is randomly sampled from a uniform distribution bound within the minimum and maximum observed range of the features.

#### Algorithm 1

COALA: Counterfactual Optimization

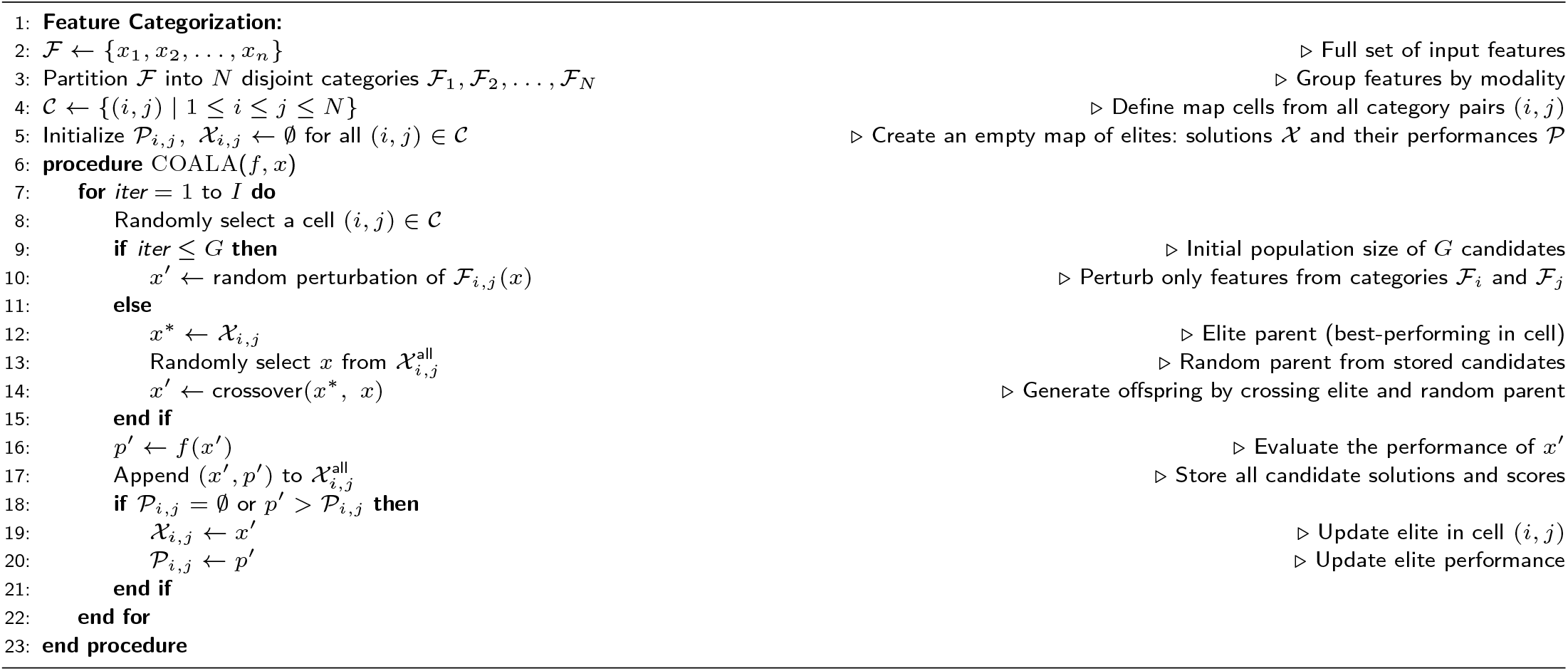

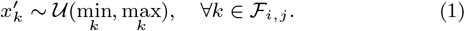

Binary features are randomly assigned values from their respective two-category sets. Repeating this process produces an initial set of candidate solutions across cells.

During the optimization phase (lines 7–8 and 12–20 of Algorithm 1), COALA continues to iteratively refine counterfactuals. In each iteration, a cell is selected at random, and two counterfactuals from that cell—one elite solution (X_*i,j*_) and one sampled at random— are selected as parents. These are recombined via crossover to produce a new candidate, which is assigned back to the same cell, regardless of whether it outperforms its parents. This promotes local exploration within the cell’s defined feature subspace. For our analyses, we used uniform crossover, but COALA also supports other crossover methods, such as single point crossover and simulated binary crossover. The optimization proceeds until a termination criterion is met, such as a maximum number of generations. This process is repeated independently for each input sample to generate optimal counterfactuals for each subject in the population.

### Actionable interpretations

For each subject, COALA will generate one counterfactual for each of *I* iterations. Each sample will have one optimal counterfactual for each cell. Every sample’s counterfactuals of a specific cell can then be analyzed with common statistical methods to identify trends in counterfactuals across the sample population. This involves clustering of the counterfactuals to identify different risk groups, then statistical tests such as ANOVA to identify risk factors that drive a sample’s risk group membership.

Because COALA searches for optimal counterfactuals within specific cells, where all features in *F*_*i,j*_ are mutable and all remaining features are held constant, differences in the resulting optimal counterfactuals across the population can be attributed to differences in the constraint features. In other words, COALA identifies the optimal values of mutable features conditional on the fixed values of the constraint features. If the relationship between features and the outcome is simple and additive, one would expect similar mutable feature changes to be optimal for all subjects. Conversely, if there is heterogeneity in the counterfactuals of the population, this suggests that the model has captured interactions between mutable and constraint features.

To identify intervention groups, subjects are clustered based on the mutable features of their optimal counterfactuals within a given cell. This reveals distinct clusters of counterfactuals, which may be candidate interventions, and the clustering process is done independently once for each cell. Although various clustering methods may be appropriate, we performed K-means clustering on the counterfactuals’ mutable feature values, which were scaled using z-score standardization. Principal component analysis (PCA) was performed on the scaled counterfactuals for visualization purposes. The optimal number of clusters was selected using the silhouette score, which quantifies the cohesion and separation of clusters.

We then identified which features characterize the different treatment clusters by performing ANOVA tests on the mutable features and computing the proportion of each feature’s variance explained by cluster membership (*η*^2^):

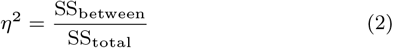

We also performed ANOVA tests on the constraint features of each cell and compute *η*^2^. All ANOVA tests were conducted on the original (unscaled) feature values to preserve interpretability. Constraint features with larger relative *η*^2^ values can be interpreted as interacting with mutable features and modulating their influence on the predicted outcome. In other words, these constraint features act as drivers in determining optimal treatments. Features were considered to substantially differentiate subgroups if *η*^2^≥0.14, corresponding to Cohen’s large effect size convention [Cohen, 1988, Richardson, 2011].

Lastly, we performed z-score standardization on the counterfactuals’ values of mutable variables and calculated the Euclidean distance between subjects. This was used to generate and visualize a similarity matrix of the study population’s optimal counterfactuals.

### SHAP analysis

We compared COALA with a SHAP-based clustering framework commonly used to identify risk subgroups in machine learning models. We apply the SHAP clustering approach found in biomedical research that uses ML models to identify actionable interventions for complex diseases [Chen et al., 2025]. SHAP values of mutable features were computed for each sample, yielding a vector of feature attributions. Subjects were then clustered with hierarchal clustering using Ward’s linkage method and Euclidean distance based on the SHAP values of mutable features to identify subgroups with similar patterns of model explanation. The number of clusters was specified based on the expected number of clusters from COALA analysis. To interpret the resulting clusters, ANOVA tests were performed on both the SHAP values and original feature values to identify which features differentiated the identified subgroups the most.

To quantitatively compare COALA and SHAP clustering, we assessed if cluster membership could be predicted from the original values of constraint features. A random forest classifier was trained on constraint feature values to predict cluster membership from COALA and SHAP and evaluated using balanced accuracy with a holdout validation set. Higher predictability indicates that the clustering structure is meaningfully organized by constraint features.

### Datasets and models

We compared COALA and SHAP-based clustering on two real biomedical datasets and a synthetic dataset, which are tabular datasets. To benchmark the COALA framework in a controlled setting, we constructed a synthetic dataset with *n* = 1000 samples, 9 input features divided into four conceptual views: genetic risk, environmental exposures, nutritional, and metabolic, and a continuous outcome *y* that represents a hypothetical health measurement (e.g., cholesterol levels) (Supplementary Table S1). The outcome was a continuous variable *y*, generated using a weighted combination of additive effects and interaction terms:

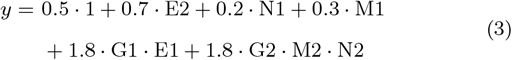

The primary biomedical dataset is derived from the National Health and Nutrition Examination Survey (NHANES), a population survey conducted by the Center for Disease Control and Prevention (CDC) [Patel et al., 2016]. NHANES was designed to estimate the prevalence of diseases, such as diabetes, in the United States. The dataset includes a range of demographic, clinical, and lifestyle variables, including sex, age, smoking history, and diet nutritional content (Supplementary Table S2). We use data from the 2017-2018 survey cycle, which integrates demographic, anthropometric, clinical, and dietary variables. The outcome variable is a binary indicator of diabetes status, defined as glycated haemoglobin (HbA1c) ≥ 6.5%, consistent with American Diabetes Association diagnostic criteria [American Diabetes Association Professional Practice Committee, 2024]. We also apply the COALA framework to a dataset derived from the Framingham Heart Study (FHS), a long-running prospective cohort study designed to identify risk factors for cardiovascular disease in adults [Dawber et al., 1951, Wilson et al., 1998]. The FHS dataset results are presented in the Supplementary Materials.

A linear regression model and a multilayer perceptron (MLP) were each trained on the synthetic dataset, and an eXtreme Gradient Boosting (XGBoost) model was trained on the real datasets, using an 80/20 train–test split. Linear regression models are limited to capturing additive relationships, while MLP and XGBoost models are capable of modeling nonlinearities and potential feature interactions [Beam and Kohane, 2018, Chen and Guestrin, 2016]. Each trained model–dataset pair was then analyzed using both the COALA and SHAP frameworks. We also analyzed a ground truth model implemented directly from the data-generating function (Equation 3) to assess whether COALA can recover known interaction patterns.

## Results

### Evaluation on synthetic datasets

We first evaluated COALA on the ground truth model for the synthetic data to investigate if COALA could identify the expected optimal counterfactual for each sample. We applied COALA by constraining synthetic genetic features and mutating environmental features. When the constraint G1 is positive, the optimal counterfactual value for E1 is positive, and is negative when G1 is negative. This is expected from the specified G1 · E1 interaction during data generation (Equation 3). We also expect that when G2 was positive, optimal counterfactual values for N2 and M2 are either both positive or both negative. When G2 is negative, one of N2 and M2 is positive, and the other negative. This pattern matches the expected results from the G2 · N2 · M2 interaction. Based on the specified interaction terms in the synthetic data, there are eight distinct optimal counterfactuals. As expected, COALA applied to the linear regression model identified identical optimal counterfactuals for all subjects, as features affect the ML prediction independently of other features. However, when applied to the ground truth model, COALA correctly identifies the expected eight distinct optimal counterfactuals (Supplementary Fig. S1A). ANOVA test of the counterfactuals’ mutable feature values revealed that the variance in mutable features E1, M2, and N2 can be explained by cluster membership (Supplementary Fig. S1B), and the variance in mutable features can be explained by the variance in the constraint feature values G1 and G2 (Supplementary Fig. S1C). Additionally, as expected E2, N1, M1, and G3 have no variation across samples when mutable, as they were not specified to have any interactions during data generation (Equation 3).

The MLP model had an MSE of 0.5431 and *R*^2^ of 0.9504 on the validation set. When applying COALA to the MLP model, G2 and G1 were identified as the constraints driving different counterfactuals as expected (Fig. 2C, E). However, the optimal value of E2, which was not specified to form interactions with other features, varied between clusters (Fig. 2B, D). There were also only six distinct clusters identified (Fig. 2A), compared to the expected eight clusters identified when COALA was applied to the ground truth model.

**Figure 2.**
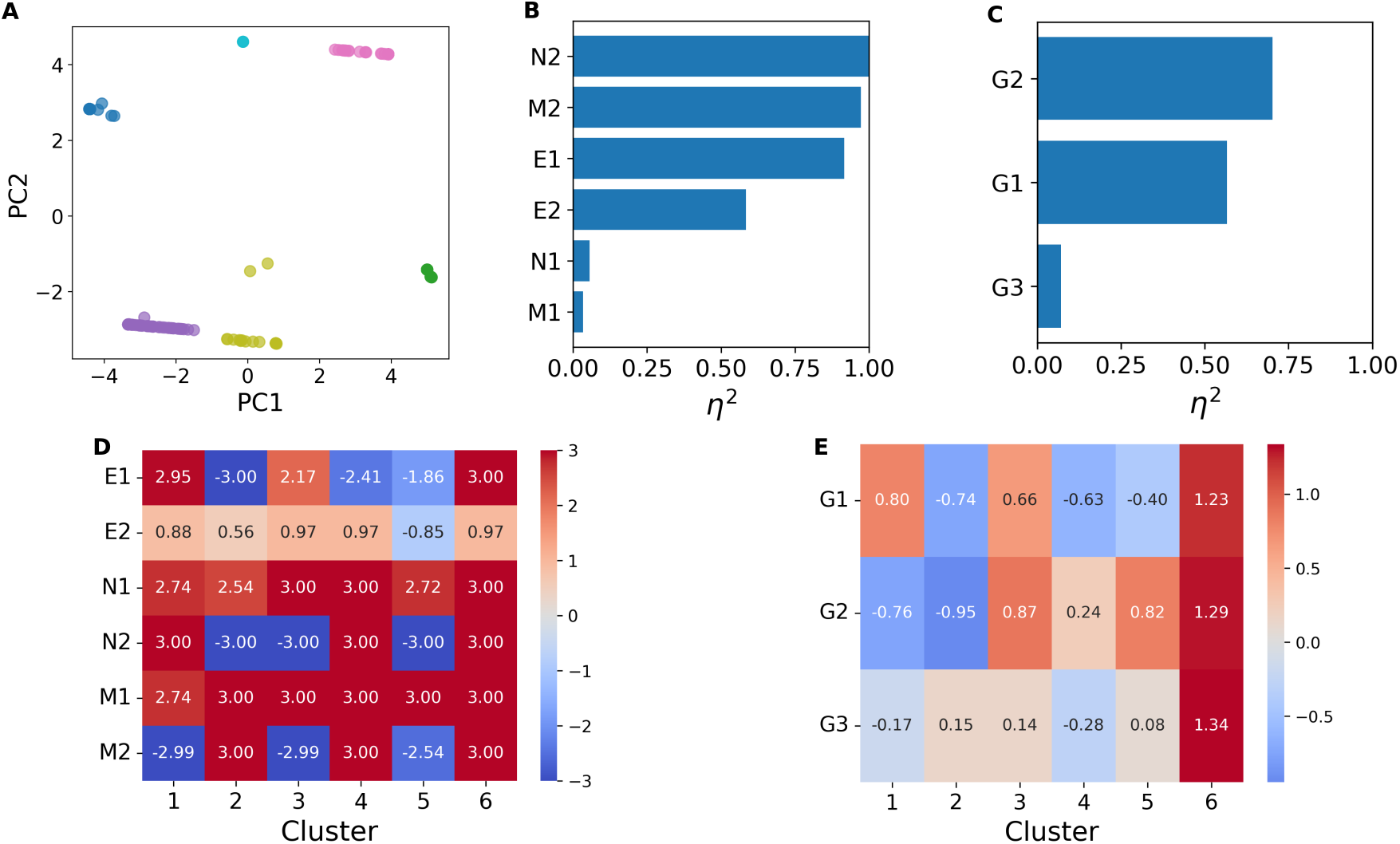
COALA analysis of the MLP model trained on the synthetic dataset. (A) PCA of mutable features across counterfactuals colored by cluster. (B) Proportion of variance in mutable features explained by cluster membership (*η*^2^). (C) Proportion of variance in constraint features explained by cluster membership (*η*^2^). (D) Mean values of mutable features by cluster. (E) Mean values of constraint features by cluster.

### Application to a real dataset

The XGBoost model trained on the NHANES dataset achieved an AUROC of 0.7541, AUPRC of 0.415, and F1 of 0.400 on the holdout validation set. The NHANES dataset serves as a proof-of-concept demonstration of COALA’s application workflow. Given the dataset’s class imbalance and the model’s limited predictive performance, COALA outputs should be interpreted as reflecting learned model behaviour rather than ground-truth clinical relationships.

After implementing COALA to identify optimal counterfactuals when only dietary features are mutable, the silhouette score identified *>* 10 clusters as the statistically optimal clustering of counterfactuals, reflecting a continuous spectrum of optimal counterfactuals rather than discrete subgroups (Fig. 3). We use K-means clustering as an approximation to partition the continuous spectrum into interpretable subgroups. Three clusters were selected to yield counterfactual subgroups with comparable sample sizes (Fig. 4A, *n*_1_=151, *n*_2_=116, *n*_3_=233). ANOVA test of subjects’ constraint features revealed waist circumference to have the largest effect size between clusters (Fig. 4C, E). Protein, total sugars, and saturated fat had the largest effect sizes among mutable features (Fig. 4B, D).

**Figure 3.**
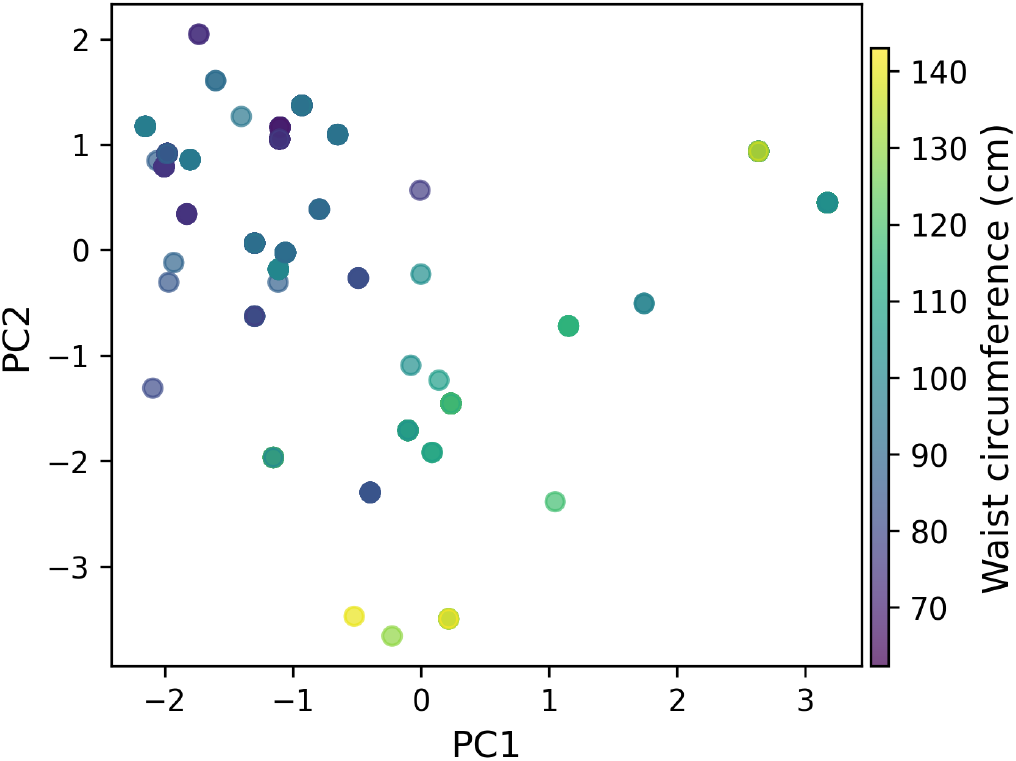
COALA analysis of the XGBoost model trained on the NHANES dataset visualized with PCA of mutable features across counterfactuals, colored by waist circumference.

**Figure 4.**
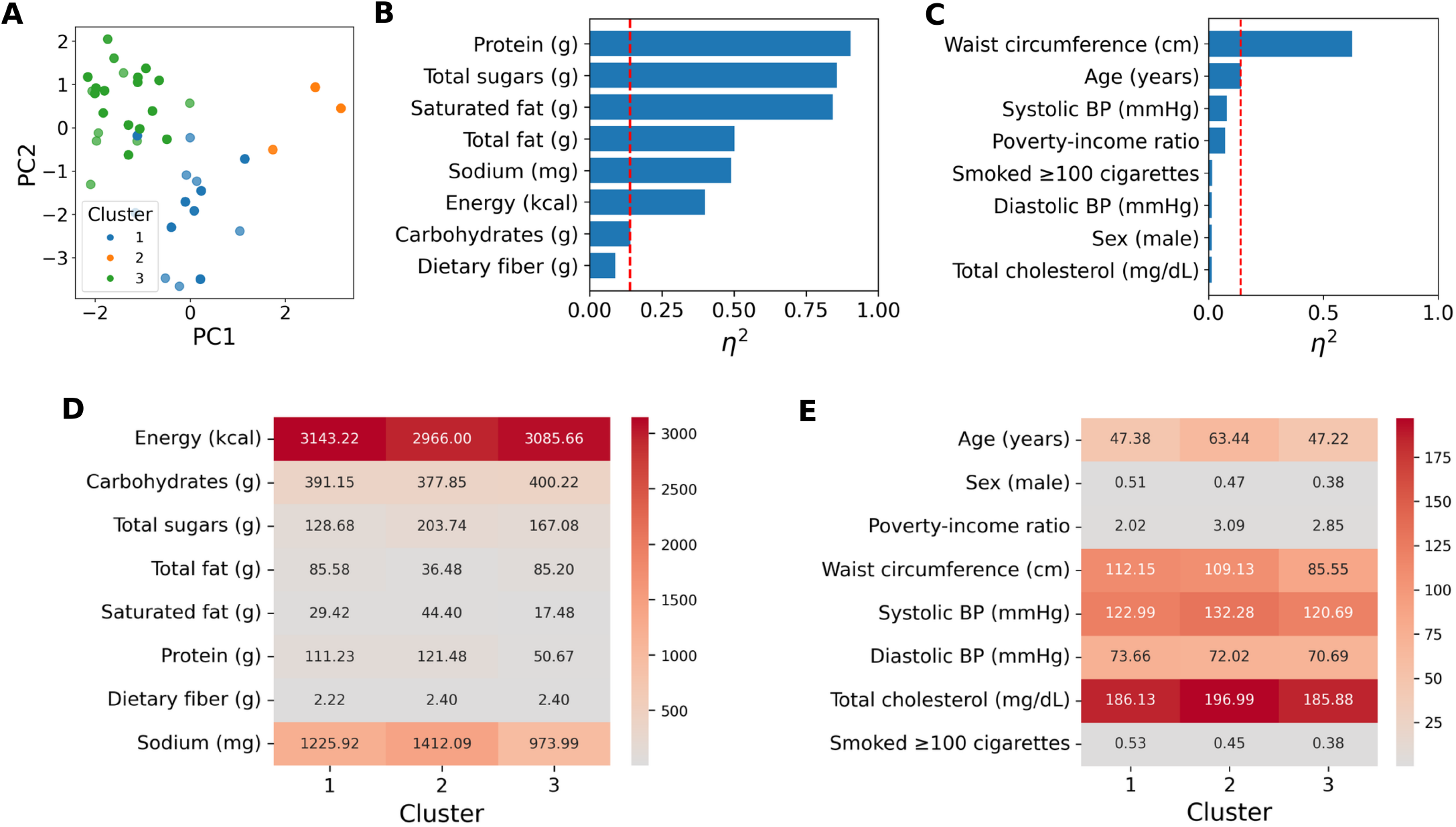
COALA analysis of the XGBoost model trained on the NHANES dataset. (A) PCA of mutable features across counterfactuals colored by cluster. (B) Proportion of variance in mutable features explained by cluster membership. (C) Proportion of variance in constraint features explained by cluster membership. The red line indicates the threshold for a large effect size (*η*^2^*≥*0.14). (D) Mean values of mutable features by cluster. (E) Mean values of constraint features by cluster.

### Population similarity network

While the overall gradient of optimal counterfactuals aligns with the waist circumference (Fig. 3), the population similarity network reveals substructures in the optimal counterfactuals that are not apparent from PCA alone (Fig. 5). There are subpopulations with similar waist circumference values but different optimal counterfactuals, suggesting other features, such as age, also meaningfully contribute to determining the optimal counterfactual (Fig. 5).

**Figure 5.**
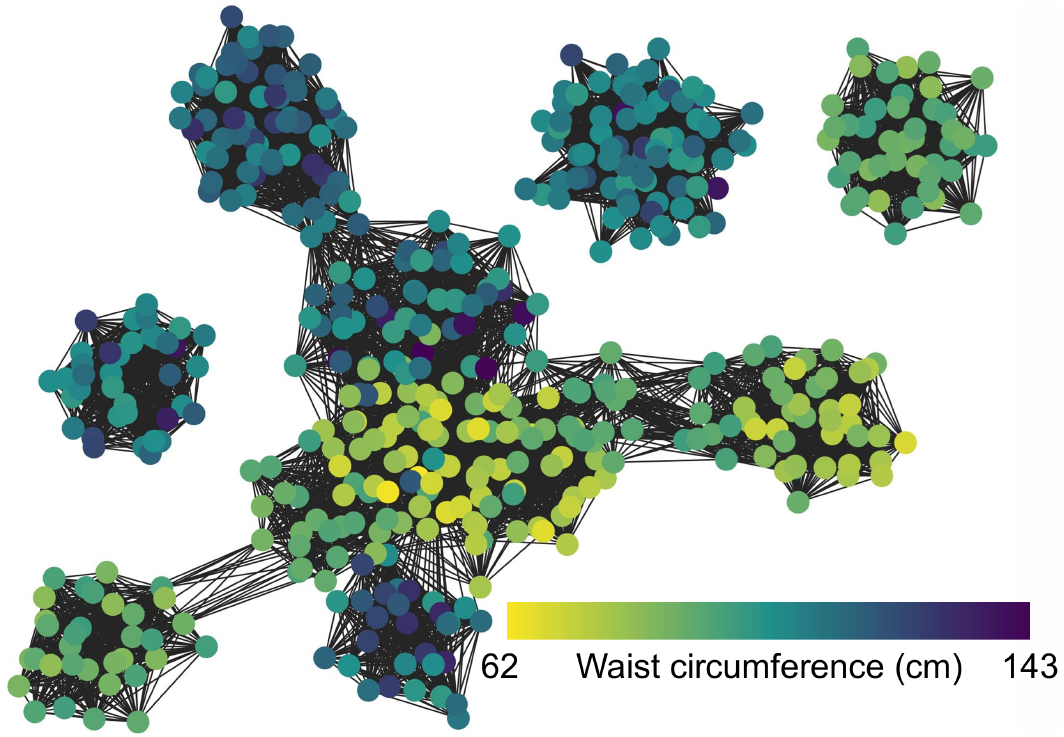
Similarity network of the population of optimal counterfactuals generated through COALA analysis on the XGBoost model trained on the NHANES dataset, colored by the subject’s waist circumference.

### Comparison with SHAP

We applied SHAP clustering to the XGBoost model and holdout NHANES dataset, grouping subjects into three clusters (Fig. 6A). SHAP clustering identified energy, dietary fiber, and carbohydrate consumption SHAP values to differentiate clusters (Fig. 6B, D). However, these three features were identified by COALA to have the smallest effect size among mutable features across counterfactuals (Fig. 4B). Features with the largest variance in SHAP values between clusters generally also have a larger variance in their original raw values (Fig. 6C, E).

**Figure 6.**
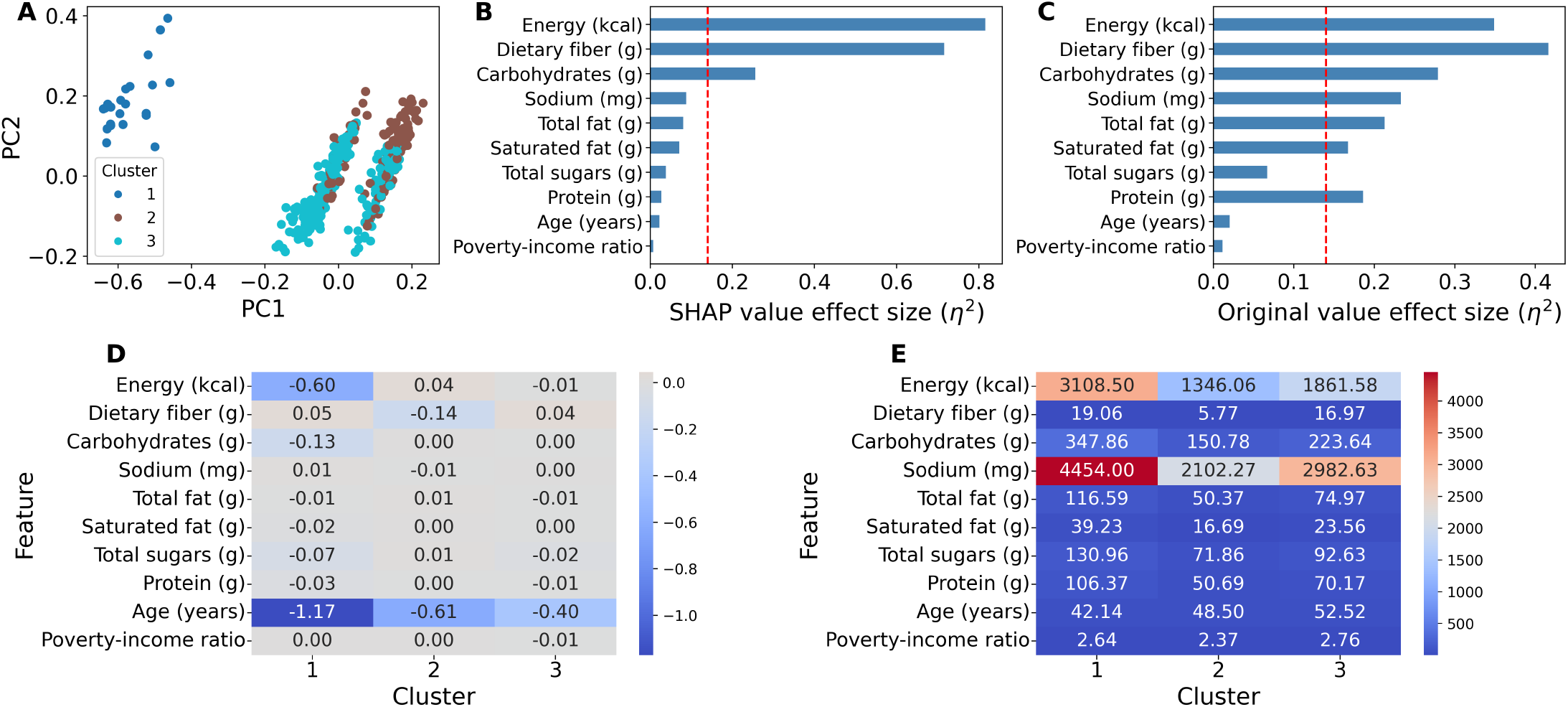
SHAP clustering analysis of the XGBoost model trained on the NHANES dataset. (A) PCA for visualization of samples’ SHAP values. (B) Proportion of variance in SHAP values explained by cluster membership for the ten features with the largest impact on prediction. (C) Proportion of variance in original feature values explained by cluster membership. The red line indicates the threshold for a large effect size (*η*^2^*≥*0.14). (D) Mean SHAP values of feature by cluster. (E) Mean original values of feature by cluster.

Constraint feature values predicted SHAP cluster membership with a balanced accuracy of 32.4%, compared to 85.4% for cluster identified with COALA (Table 1). In both counterfactual and SHAP value clustering, only the mutable features were used to assign cluster membership, but COALA clusters are more organized by constraint features. COALA demonstrated even stronger constraint feature predictability in the Framingham Heart Study dataset, achieving 100% holdout accuracy compared to 53.3% for SHAP-based clustering (Supplementary Table S4), further supporting the generalizability of the framework across biomedical datasets.

**Table 1.**
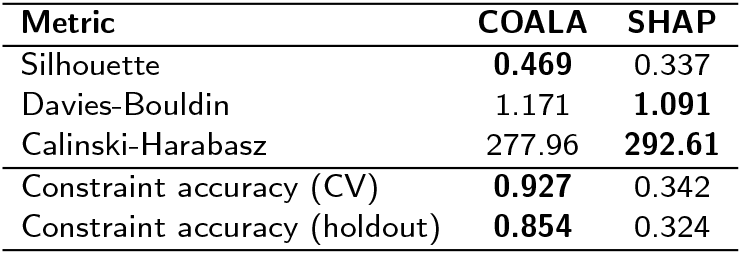
Comparison of clustering quality metrics between COALA and SHAP-based clustering on the NHANES dataset. Bold indicates the better value for each metric. With the exception of the Davies-Bouldin score, a higher score indicates better clustering.

## Discussion

COALA was applied to a ground truth model of the synthetic data to confirm that the expected optimal counterfactuals can be identified. As expected, each of the eight distinct optimal counterfactuals only differ in features that form interactions with genetic factors, G1 and G2 (Supplementary Fig. S1). Similarly, G1 and G2 as constraints have large effect sizes between groups (*η*^2^ *>* 0.6), suggesting that they dictate the optimal values of synthetic actionable features (E1, M2, and N2). However, when COALA was applied to an MLP model, E2 was incorrectly assigned a large effect size, where low E2 is beneficial to cluster five subjects (Fig. 2B, D). Thus, COALA can be used to assess whether an ML model has accurately learned the underlying feature interactions in the data. Here, COALA reveals that this MLP model would be less generalizable for subjects in cluster 4, who have moderately low G1 and high G2 (Fig. 2C, E).

When applied to the NHANES dataset, COALA revealed waist circumference to be the main determinant of optimal dietary counterfactuals (Fig. 3, 4). This is consistent with the established role of waist circumference as a strong predictor of diabetes risk [Klein et al., 2007]. COALA’s results are also consistent with evidence that related physiological factors, such as BMI, predict glycemic responses and can be used to prescribe personalized diets [Zeevi et al., 2015]. Constraint features can predict counterfactual cluster membership with high accuracy (Table 1), but the similarity network also reveals that the relationship between constraints and optimal counterfactuals can involve complex, nonlinear structure (Fig. 5), motivating future work on more nuanced approaches to characterizing this relationship.

SHAP-based clustering identifies groups of subjects who share similar features that impacted the prediction, grouping together subjects with similar feature attributions but SHAP does not reveal if subjects would respond similarly when features are changed (Fig. 6, Table 1). SHAP is able to reveal features that drive the current ML prediction (Fig. 6). However, it is not able to capture how constraint features modulate the effect mutable features have on the predicted outcome, as constraint features are not able to predict SHAP clustering membership (Table 1).

In contrast, how COALA identifies subgroups is more closely aligned with personalized intervention strategies. COALA identified risk groups by examining how certain features modulate the influence of other features on the outcome, revealing interaction effects (Fig. 4B, C). Thus, constraint features can predict the counterfactual cluster membership of a subject (Table 1). Biological outcomes are often shaped by interactions among factors, where the effect of one factor depends on the context set by others. SHAP assesses an instance retrospectively, and while it may provide insight into what changes could be made, how beneficial it would be to improving the predicted outcome as a personalized intervention is not clear. COALA is designed to assess instances prospectively, focusing on the intervention, and reveals how specific changes would impact the predicted outcome differently across subjects. Using SHAP clustering to group subjects in a population can be described as finding risk groups, but using COALA may be described as finding groups for targeted treatment or intervention.

Counterfactuals by nature imply causality, but it is important to note that the causality directly applies only to the ML model, and not to the true biological effects, as such causal inferences require causal models [Molnar, 2025]. However, causal models often struggle with more complex patterns, such as nonlinear interactions and context-specific effects [Runge et al., 2019]. In general, causal modeling requires strong domain knowledge of the features and outcome to make assumptions for causal modeling, which can be challenging with a high dimensional dataset for a complex outcome [Berkessa et al., 2025]. Even though COALA can not fully validate if trends identified by an ML model are truly causal, it can help to generate new hypotheses and inform users of ways to build upon a causal model.

Because COALA is an evolutionary algorithm, computational costs increase linearly with sample size. Future work could explore strategies to decrease the computational resources required to search for the optimal counterfactual. Additionally, a limitation of COALA is that counterfactuals are generated without enforcing real-world feasibility constraints beyond the observed range of feature values [Mothilal et al., 2020]. As a result, optimal counterfactuals may not always be realistically achievable, such as counterfactuals that require a drastic lifestyle change or counterfactuals that are inconsistent with health recommendations. For example, COALA identified optimal counterfactuals with high energy intake *>*3000kcal. Future implementations could incorporate domain knowledge and feasibility constraints to generate more realistic counterfactuals. Furthermore, COALA interprets the learned behaviour of the ML model rather than ground-truth biological relationships, meaning outputs are only as reliable as the underlying model [Molnar, 2025]. However, this also enables COALA to serve as a model validation tool, revealing cases where a model has not accurately learned expected feature interactions.

While COALA was initially designed and applied to ML models trained on tabular data, COALA’s approach of using counterfactuals is not limited to specific types of dataset. COALA is also model agnostic, as it treats all models as a black box. Only the prediction function is needed to optimize for counterfactuals. For interpretable models, such as linear regression and K-nearest neighbor (KNN) models, that already inherently provide transparent explanations for users to generate counterfactuals on their own, COALA is not necessary. However, application to black box models, such as MLPs and gradient boosted trees in this paper, is when COALA can thrive. COALA addresses a fundamental gap in ML model interpretations that lead to actionable insights. Existing XAI methods identify which features influence a prediction, but cannot reveal how fixed biological and clinical factors determine what intervention is optimal for a given subject. COALA bridges this gap by optimizing for counterfactuals at a population level, and revealing which features drive variations in optimal counterfactuals across subjects. As ML models become increasingly integrated into biomedical research, COALA is a step in the direction of leveraging ML models to make actionable interpretations.

## Author contributions statement

Bryant Han (Conceptualization [lead], Formal analysis [lead], Methodology [lead], Validation [lead], Visualization [lead], Writing-original draft [lead]), Qingling Duan (Conceptualization [supporting], Methodology [supporting], Writing-review & editing [supporting], Ting Hu (Conceptualization [lead], Methodology [lead], Writing-review & editing [supporting])

## Acknowledgments

The authors used Claude (Anthropic) for assistance with code development. All content was reviewed and verified by the authors, who take full responsibility for the integrity of the work.

