## Supplementary Data for "Identifying intervention strategies from machine learning models with COALA: a counterfactual optimization framework"

This PDF contains Supplementary Tables and Figures referenced in the main text.

#### Contents

**1 Supplementary Tables** **2**

**2 Supplementary Figures** **4**

**3 Framingham dataset analysis** **6**

**4 Multi-cell analyses** **11**

### 1 Supplementary Tables

**Table S1:** Categorization of features in the synthetic dataset when COALA is used in multicell mode and distributions of features.

| **Category** | **Feature Description** |
|---|---|
| Genetics | G1 $\sim \mathcal{N}(0,1)$ |
| | G2 $\sim \mathcal{N}(0,1)$ |
| | G3 $\sim \mathcal{N}(0,1)$ |
| Environment | E1 $\sim \mathcal{N}(0,1)$ |
| | E2: binary, $p = 0.2$ |
| Nutritional | N1 $\sim \mathcal{N}(0,1)$ |
| | N2 $\sim \mathcal{N}(0,1)$ |
| Metabolic | M1 $\sim \mathcal{N}(0,1)$ |
| | M2 $\sim \mathcal{N}(0,1)$ |

**Table S2:** Description of outcome and features included in the NHANES 2017–2018 analysis.

| Variable | Category | Description |
|---|---|---|
| Diabetes status (Outcome) | Outcome | Binary indicator of diabetes status, defined as glycated hemoglobin (HbA1c) $\geq 6.5\%$, consistent with American Diabetes Association diagnostic criteria. |
| Age (years) | Constrained | Age of the participant at time of survey, measured in years. |
| Sex (male) | Constrained | Biological sex of the participant, coded as 1 for male and 0 for female. |
| Poverty-income ratio | Constrained | Ratio of family income to the federal poverty level, used as a proxy for socioeconomic status. |
| Waist circumference (cm) | Constrained | Waist circumference measured at the level of the iliac crest, in centimetres. |
| Systolic BP (mmHg) | Constrained | Systolic blood pressure measured at examination, in millimetres of mercury. |
| Diastolic BP (mmHg) | Constrained | Diastolic blood pressure measured at examination, in millimetres of mercury. |
| Total cholesterol (mg/dL) | Constrained | Total serum cholesterol concentration measured at examination (mg/dL). |
| Smoked $\geq$100 cigarettes | Constrained | Indicator variable for whether the participant reported smoking at least 100 cigarettes in their lifetime. |
| Energy (kcal) | Mutable (dietary) | Total energy intake reported from a 24-hour dietary recall, measured in kilocalories. |
| Carbohydrates (g) | Mutable (dietary) | Total carbohydrate intake reported from a 24-hour dietary recall, measured in grams. |
| Total sugars (g) | Mutable (dietary) | Total sugar intake reported from a 24-hour dietary recall, measured in grams. |
| Total fat (g) | Mutable (dietary) | Total fat intake reported from a 24-hour dietary recall, measured in grams. |
| Saturated fat (g) | Mutable (dietary) | Saturated fat intake reported from a 24-hour dietary recall, measured in grams. |
| Protein (g) | Mutable (dietary) | Total protein intake reported from a 24-hour dietary recall, measured in grams. |
| Dietary fiber (g) | Mutable (dietary) | Total dietary fiber intake reported from a 24-hour dietary recall, measured in grams. |
| Sodium (mg) | Mutable (dietary) | Total sodium intake reported from a 24-hour dietary recall, measured in milligrams. |

### 2 Supplementary Figures

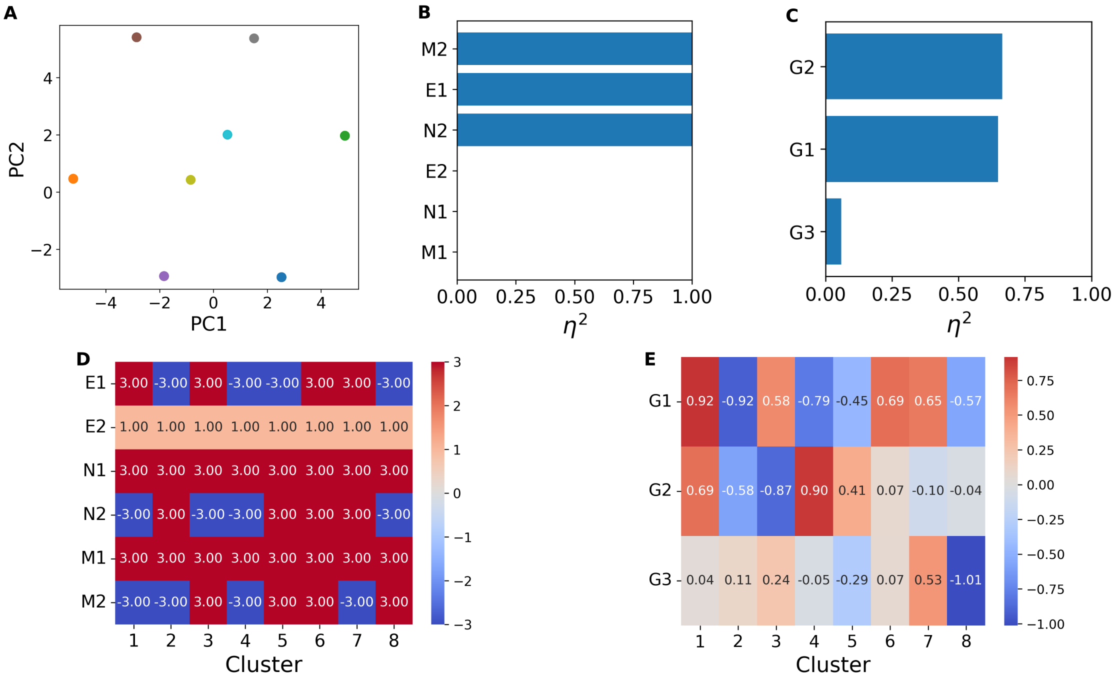


**Figure S1:** COALA analysis of the ground truth model for the synthetic dataset. (A) PCA of mutable features across counterfactuals colored by cluster. (B) Proportion of variance in mutable features explained by cluster membership ($\eta^2$). (C) Proportion of variance in constraint features explained by cluster membership ($\eta^2$). (D) Mean values of mutable features by cluster. (E) Mean values of constraint features by cluster.

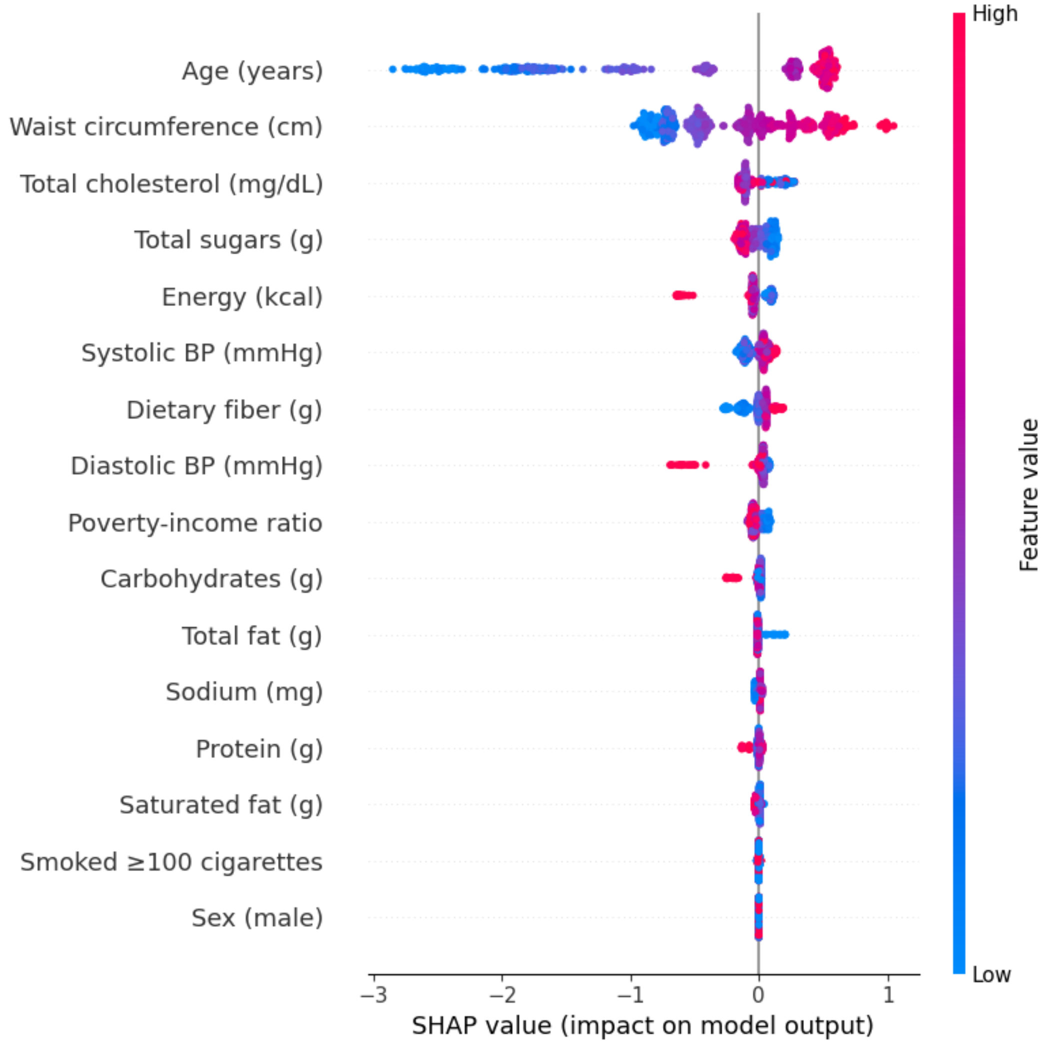


**Figure S2:** SHAP value distributions of features used in the XGBoost model's predictions on the NHANES dataset.

### 3 Framingham dataset analysis

**Table S3:** Description of outcome and features included in the Framingham Heart Study analysis.

| **Variable** | **Description** |
|---|---|
| Ten-year coronary heart disease (Outcome) | Binary indicator of whether the participant experienced a coronary heart disease (CHD) event within 10 years of baseline examination, as adjudicated in the Framingham Heart Study. |
| Sex (male) | Biological sex of the participant, coded as 1 for male and 0 for female. |
| Age (years) | Age of the participant at baseline examination, measured in years. |
| Current smoker | Indicator variable denoting whether the participant was an active smoker at baseline. |
| Cigarettes per day | Average number of cigarettes smoked per day among current smokers. |
| BP medication | Indicator variable denoting whether the participant was receiving antihypertensive medication at baseline. |
| Stroke history | Indicator variable for a prior history of stroke before baseline examination. |
| Hypertension | Indicator variable for the presence of hypertension at baseline, defined using standard clinical criteria and/or antihypertensive medication use. |
| Diabetes | Indicator variable for baseline diabetes status, based on clinical diagnosis or laboratory criteria. |
| Total cholesterol | Total serum cholesterol concentration measured at baseline examination (mg/dL). |
| Systolic blood pressure | Systolic blood pressure measured at baseline examination (mmHg). |
| Diastolic blood pressure | Diastolic blood pressure measured at baseline examination (mmHg). |
| Body mass index (BMI) | Body mass index calculated as weight in kilograms divided by height in meters squared (kg/m$^2$). |
| Resting heart rate | Resting heart rate measured at baseline examination (beats per minute). |
| Blood glucose | Fasting or random blood glucose concentration measured at baseline examination (mg/dL). |

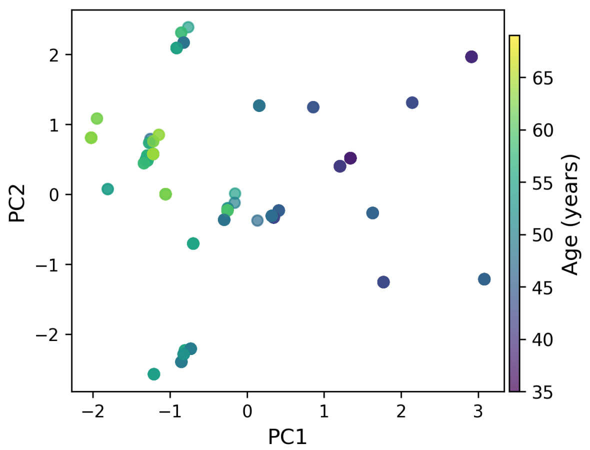


**Figure S3:** COALA analysis of the XGBoost model trained on the FHS dataset visualized with PCA of mutable features across counterfactuals, colored by age.

**Table S4:** Comparison of clustering quality metrics between COALA and SHAP-based clustering on the FHS dataset. Bold indicates the better value for each metric. With the exception of the Davies-Bouldin score, a higher score indicates better clustering.

| **Metric** | **COALA** | **SHAP** |
|---|---|---|
| Silhouette | 0.370 | **0.386** |
| Davies-Bouldin | 1.024 | **0.879** |
| Calinski-Harabasz | 146.20 | **213.41** |
| Constraint accuracy (CV) | **0.968** | 0.487 |
| Constraint accuracy (holdout) | **1.000** | 0.533 |

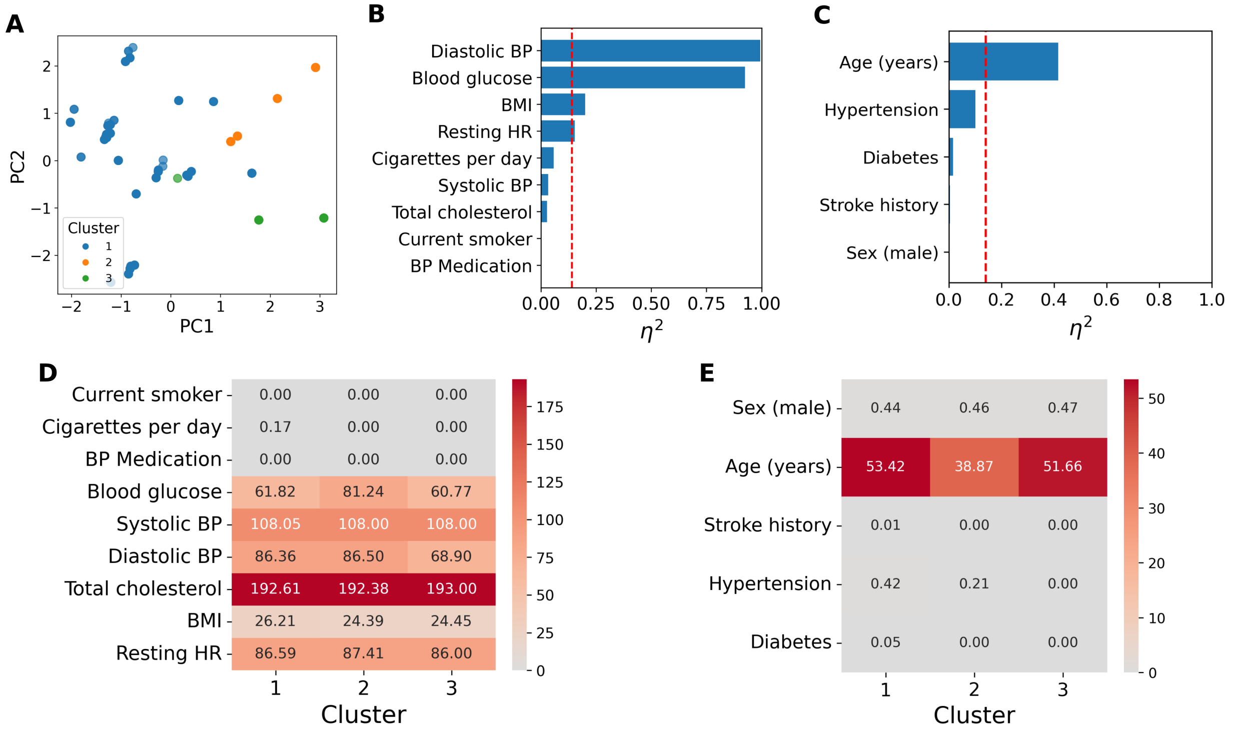


**Figure S4:** COALA analysis of the XGBoost model trained on the FHS dataset. (A) PCA of mutable features across counterfactuals colored by cluster. (B) Proportion of variance in mutable features explained by cluster membership. (C) Proportion of variance in constraint features explained by cluster membership. The red line indicates the threshold for a large effect size ($\eta^2 \geq 0.14$). (D) Mean values of mutable features by cluster. (E) Mean values of constraint features by cluster.

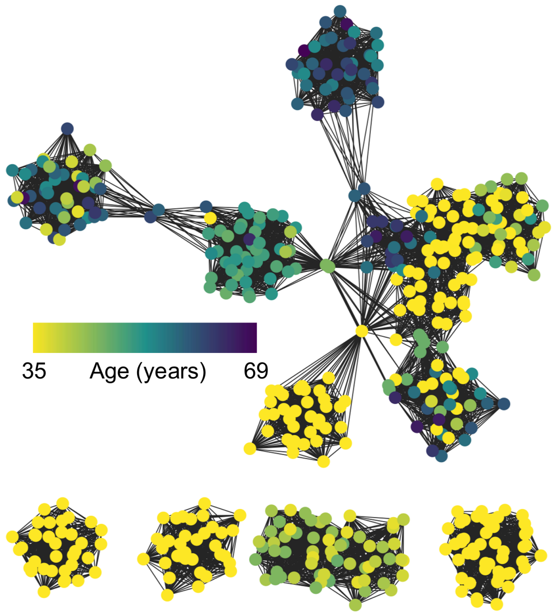


**Figure S5:** Similarity network of the population of optimal counterfactuals generated through COALA analysis on the XGBoost model trained on the FHS dataset, colored by the subject's age.

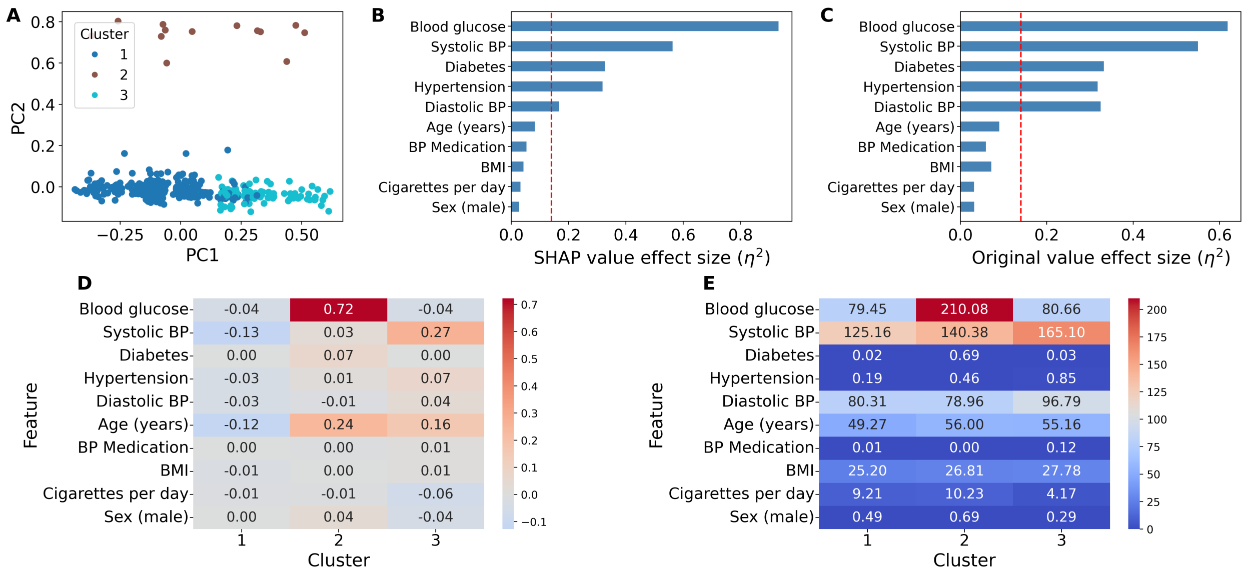


**Figure S6:** SHAP clustering analysis of the XGBoost model trained on the FHS dataset. (A) PCA for visualization of samples' SHAP values. (B) Proportion of variance in SHAP values explained by cluster membership for the ten features with the largest impact on prediction. (C) Proportion of variance in original feature values explained by cluster membership. The red line indicates the threshold for a large effect size ($\eta^2 \geq 0.14$). (D) Mean SHAP values of feature by cluster. (E) Mean original values of feature by cluster.

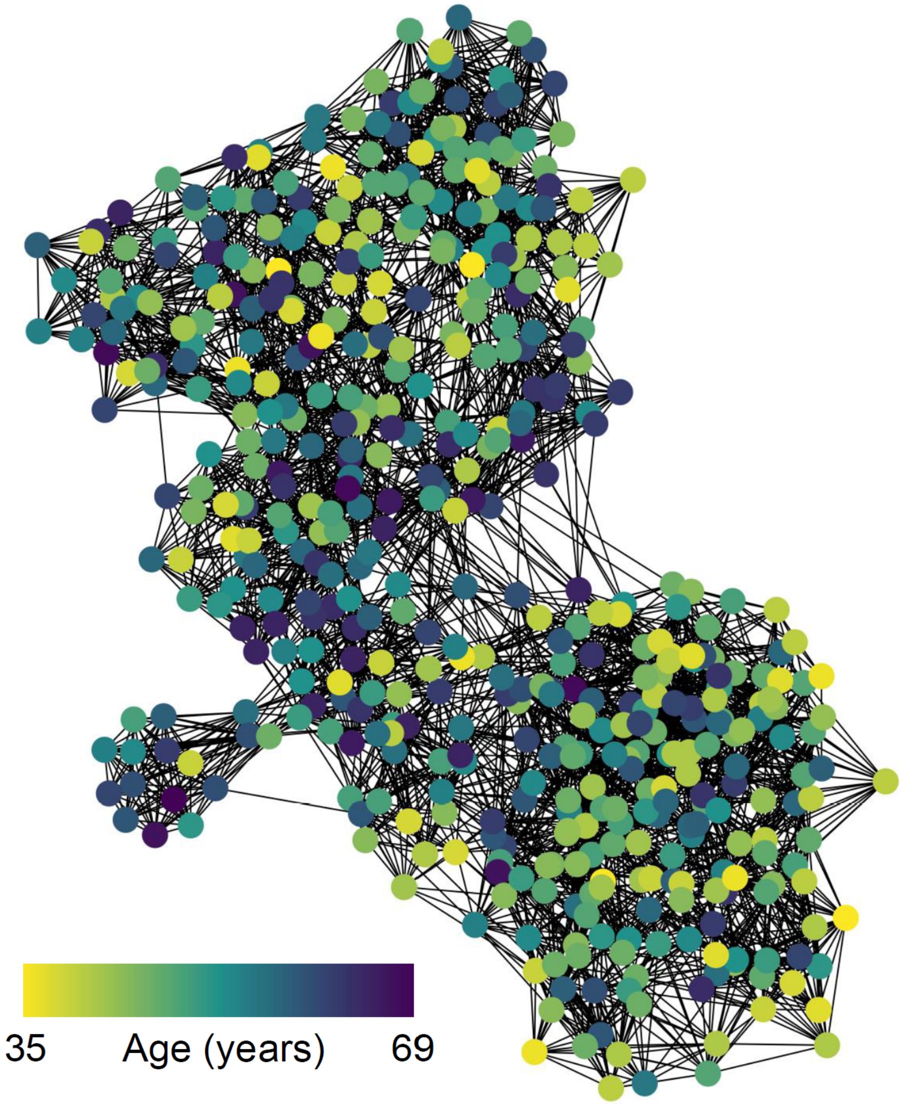


**Figure S7:** Similarity network of the population of SHAP matrices generated for test samples, from the XGBoost model and FHS dataset. Nodes are colored by age.

### 4 Multi-cell analyses

COALA analysis with multiple cells allowed for the identification of specific feature pairs that interact with each other in the synthetic dataset, such as identify E1 as a mutable that is driven by G1 when only environment features are mutable (Figure S8). There were cells where no constraint was identified to reveal the G2 · N2 · M2 interaction (Figure S8C), because although there are two clusters of optimal counterfactuals for M2 (Figure S8B), different groups can share the same counterfactual (e.g., negative G2 and N2 benefit from the same change as positive G2 and N2).

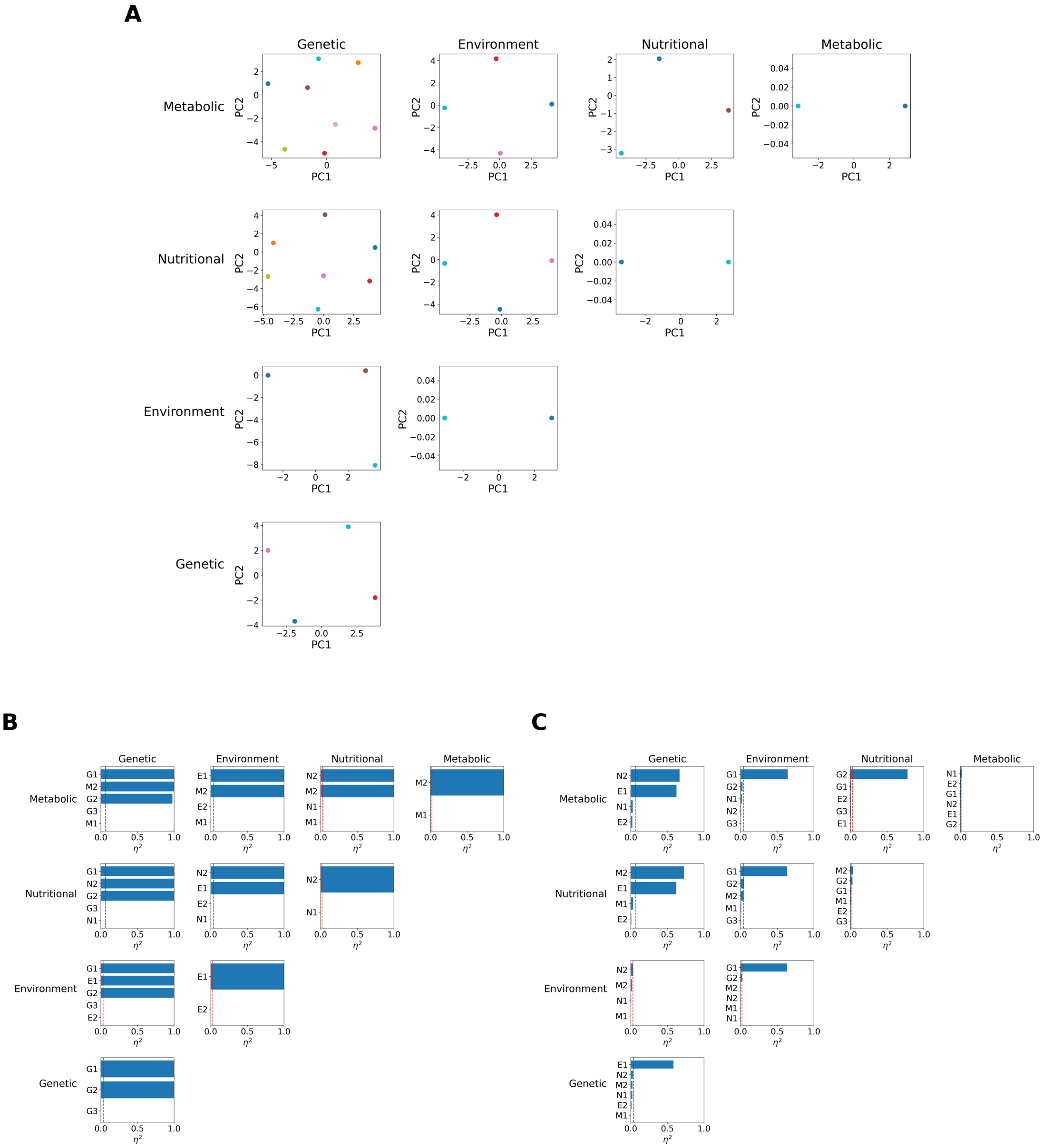


**Figure S8:** COALA analysis of ground truth model for the synthetic dataset with multiple cells specified, where subplot shows results for when certain feature categories are mutable (E.g., top left subplot shows results for counterfactuals when genetic and metabolic features are mutable). (A) PCA of mutable features across counterfactuals colored by cluster membership. (B) Proportion of variance in mutable features explained by cluster membership ($\eta^2$). (C) Proportion of variance in constraint features explained by cluster membership ($\eta^2$).

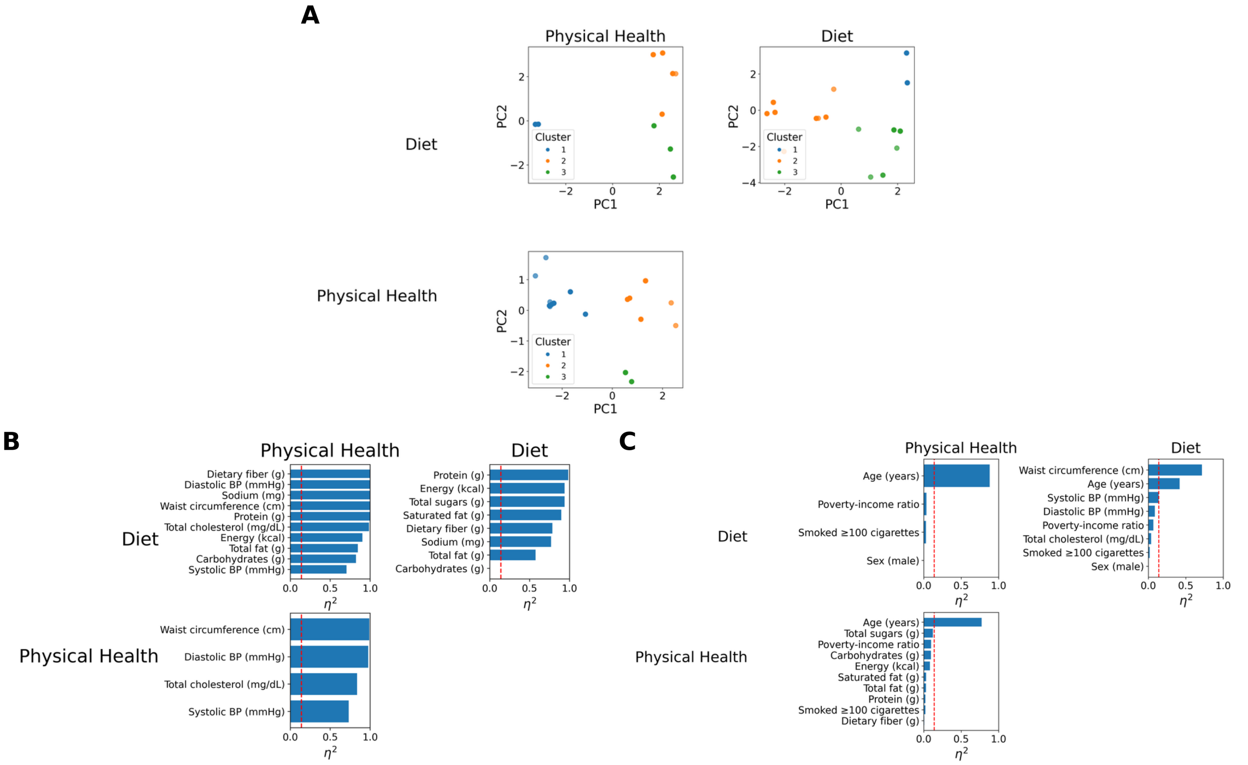


**Figure S9:** Grid of PCAs of optimal counterfactuals from COALA analysis of XGBoost model's performance on the NHANES dataset in multicell mode. Each subplot shows the variance in counterfactuals when different feature categories are mutable (E.g., top left PCA shows variance in counterfactuals where smoking and medication variables are mutable). Points are colored by cluster membership. (A) PCA of mutable features across counterfactuals colored by cluster membership. (B) Proportion of variance in mutable features explained by cluster membership ($\eta^2$). (C) Proportion of variance in constraint features explained by cluster membership ($\eta^2$).
